# Structural basis of CBP/p300 recruitment by the microphthalmia-associated transcription factor

**DOI:** 10.1101/2022.08.28.505448

**Authors:** Alexandra D. Brown, Kathleen L. Vergunst, Makenzie Branch, Connor M. Blair, Denis J. Dupré, George S. Baillie, David N. Langelaan

## Abstract

The microphthalmia-associated transcription factor (MITF) is a master regulator of the melanocyte cell lineage. Aberrant MITF activity can lead to multiple malignancies including skin cancer, where it modulates the proliferation and invasiveness of melanoma. MITF-dependent gene expression requires recruitment of transcriptional co-activators such as CBP/p300, but details of this process are not fully defined. Here, we investigate the structural and functional interaction between the MITF N-terminal transactivation domain (MITF_TAD_) and CBP/p300. A combination of pulldown assays and nuclear magnetic resonance spectroscopy determined that MITF binds both TAZ1 and TAZ2 domains of CBP/p300 with high affinity. The solution-state structure of the MITF_TAD_:TAZ2 complex reveals that MITF interacts with a hydrophobic surface of TAZ2, while remaining relatively dynamic. Peptide array and mutagenesis experiments determined that an acidic motif is integral to the MITF_TAD_:TAZ2 interaction and is necessary for transcriptional activity of MITF. Peptides that bind to the same surface of TAZ2 as MITF_TAD_, such as the adenoviral protein E1A, are capable of displacing MITF from TAZ2 and inhibiting transactivation. These results provide mechanistic insight into co-activator recruitment by MITF that are fundamental to our understanding of MITF targeted gene regulation and melanoma biology.

## INTRODUCTION

The microphthalmia family of transcription factors (MiT/TFE) is comprised of four closely related members, the microphthalmia-associated transcription factor (MITF), transcription factor EB (TFEB), TFE3, and TFEC (1). MITF is a master transcriptional regulator of melanocytes, the pigment producing cells of the skin (2), where it is essential for development and differentiation by controlling expression of many genes involved in growth, metabolism, and cell survival (3). Mutation to MITF can cause the failure of melanocytes to form, resulting in genetic diseases such as Waardenburg Syndrome type 2 and Tietz syndrome, which are characterized by skin, eye, and hearing defects (4, 5). Aberrant MITF activity has also been linked to the melanocyte-derived skin cancer, melanoma, where MITF is critical in modulating the proliferation and invasiveness of the disease (6, 7). MITF is a recognized lineage-specific oncogene in melanoma where it has a prosurvival role (3, 6, 9). While overexpression of MITF drives melanoma progression, long-term MITF suppression can lead to cellular senescence (7).

MiT/TFE family members are similar in structure and composed of a characteristic set of domains. The central basic helix-loop-helix (bHLH) leucine zipper domain is highly homologous and responsible for dimerization and DNA-recognition of canonical E-box sequences (CANNTG) in gene promoter regions (10). MiT/TFE family members also share a transactivation domain (TAD) that is N-terminal to the bHLH (Figure 1A). While divergent in sequence, these TADs allow MITF and other MiT/TFE members to recruit protein co-activators for enhanced transcription of targeted genes (11). Co-factor recruitment by activation domains is often facilitated by small recognition motifs, including the ΦXXΦΦ motif (with Φ representing a hydrophobic residue and X any amino acid), which is conserved in the MiT/TFE family (Figure 1B) and also present in other transcription factors such as E2A, HIF-1α, p53, STAT1 and STAT2 (12–16).

**Figure 1.**
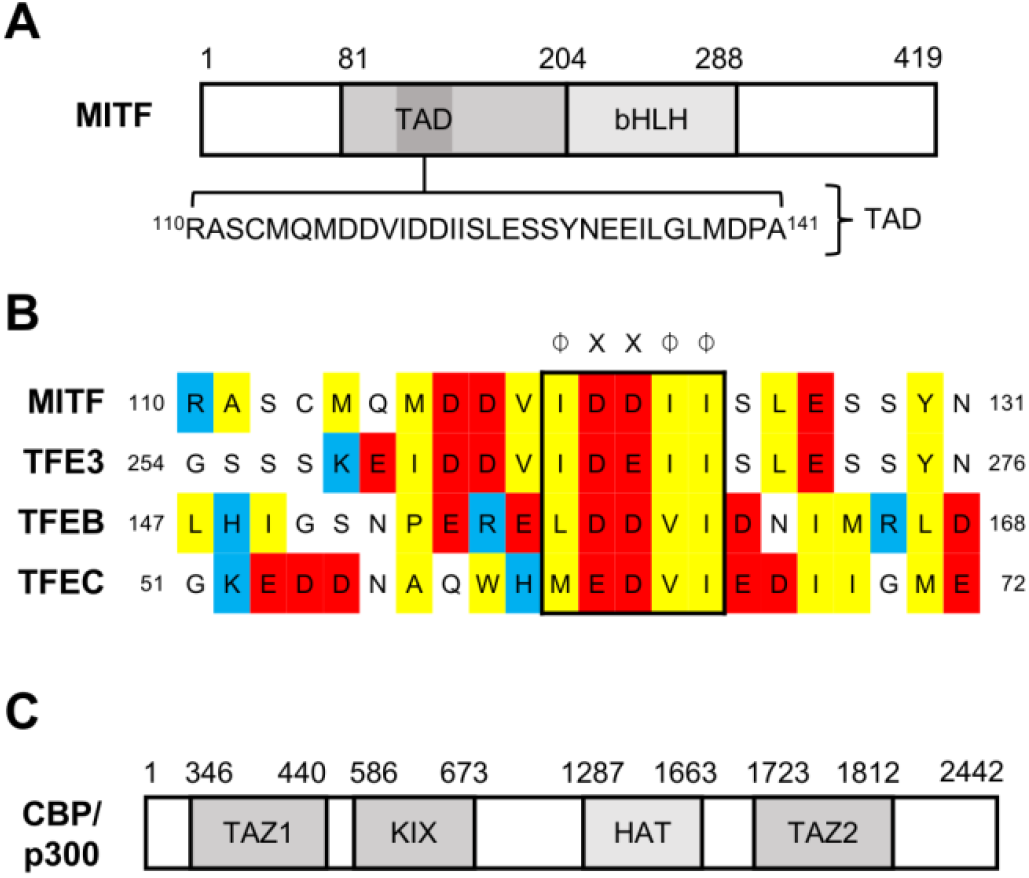
Domain architecture of MITF and CBP/p300. **(A)** Schematic of MITF illustrating the acidic N-terminal transactivation domain (TAD) and basic helix-loop-helix leucine zipper DNA binding domain (bHLH), numbered residues indicate domain boundaries. **(B)** Alignment of ΦXXΦΦ containing sequences conserved amongst activation domains of MITF, TFE3, TFEB, and TFEC, where Φ represents a hydrophobic amino acid and X any amino acid. **(C)** Domains of CBP/p300 including the catalytic histone acetyltransferase domain (HAT), and protein-interacting domains KIX, TAZ1, and TAZ2, numbering is in accordance with native protein sequence of p300.

Interacting partners with the MITF_TAD_ most notably includes the histone acetyltransferase cAMP-response element-binding protein (CREB)-binding protein (CBP), and its close homolog E1A-binding protein (p300) (17). CBP/p300 are key histone acetyltransferases that accumulate at the promotors of over 16,000 human genes (18) and interact with over 400 known binding partners (19), making them among the most heavily connected nodes in the mammalian interactome. CBP/p300 is a large multimodular protein, with folded domains including a histone acetyltransferase (HAT) domain and multiple protein-binding domains that are separated by intrinsically disordered regions (Figure 1C). The catalytic HAT domain promotes the pre-initiation complex of transcription through its acetylation of relevant histones, thereby loosening the chromatin and improving the availability of the associated DNA to be transcribed (20). The transcription adaptor zinc finger (TAZ1/2) and kinase inducible domains (KIX) form interaction sites for intrinsically disordered activation sites of transcription factors, and thus act as a conduit between a variety of transcription factors and transcriptional machinery (21, 22). Transcription factors complete for limited quantities of CBP/p300 (23), and some viral oncoproteins, such as adenovirus early region 1 A (E1A), deregulate the hosts’ cell cycle by sequestering CBP/p300 through tight binding interactions with the TAZ2 domain (24, 25).

Despite the importance of MITF for melanocyte differentiation and melanoma biology, the mechanistic details of how MITF interacts with CBP/p300 to induce MITF-target gene expression are not fully understood. Here, we characterize the structure and function of the MITF_TAD_ and its interactions with different domains of CBP/p300 using a combination of structural biology, pulldown, and transactivation assays. We then evaluate the ability of an E1A-derived peptide to inhibit TAZ2 recruitment by MITF and impede MITF transcriptional activity.

## MATERIALS AND METHODS

### Plasmid preparation

Full length MITF isoform M cDNA was purchased from ThermoFisher Inc. (Genbank Accession no. BC065243.1). Residues 81-204, 110-161, and 110-141 of MITF-M were amplified by PCR and subcloned into a modified pET21 vector containing upstream sequences coding for a hexahistidine tag (His_6_), the B1 domain of *Streptococcus* protein G (GB1), and a tobacco etch virus protease cleavage site to create pGB1-MITF_81-204_, pGB1-MITF_110-161_, and pGB1-MITF_TAD_ respectively. Residues 1-204, 81-204, 1-100, and 289-419 of MITF were cloned into pCMV-GAL4 vector gifted by Liqun Luo (Addgene plasmid # 24345) for use in transcriptional activation assays (pGAL4-MITF) (26). Full-length human p300 (residues 1-2414) was subcloned into pcDNA3.1(+) mammalian expression vector (pcDNA-p300) for transfection. Plasmids coding for TAZ1 (residues 346–440 of CBP), TAZ2 (residues 1723-1812 of p300 with four stabilizing mutations C1738A, C1746A, C1789A, C1790A), and KIX (residues 586–673 of CBP) were provided by Dr. Steven Smith (Queens University, Kingston, ON) (12). A pET21 derived plasmid containing sequences encoding His_6_, GB1, TEV, and residues 54-82 (CR1) from adenovirus early region 1 A (pGB1-E1A_CR1_) was synthesized by BioBasic Inc, and the E1A_CR1_ sequence was then subcloned into pcDNA3-GFP gifted by Doug Golenbock (Addgene plasmid # 13031) (pGFP-E1A_CR1_). MITF deletion mutants (ΔTAD, Δ117-124, Δ123-127) were generated from pGB1-MITF_81-204_ and pGAL4-MITF_1-204_ using the Q5® QuickChange site-directed mutagenesis kit (New England Biolabs) according to the manufacturer’s protocol. The fidelity of all constructs was confirmed by DNA sequencing.

### Protein expression and purification

Plasmids pGB1-MITF, pGB1-TAZ2, and pGB1-E1A_CR1_ were transformed into chemically competent *E. coli* BL21 (DE3) cells for recombinant protein expression. Cultures were grown in LB or ^15^N-or ^15^N/^13^C enriched M9 Minimal Media (27) at 37°C to an optical density at 600 nm of 0.6-0.8, following which recombinant protein expression was induced with the addition of isopropyl 1-thio-β-D-galactopyranoside (IPTG; 0.5 mM). Cultures were incubated for 4 h at 37°C or overnight at 20°C and then harvested by centrifugation. Purification of TAZ1 and KIX occurred as previously described (12). For purification of GB1-TAZ2, cell pellets were resuspended in 30 mL of denaturing lysis buffer (20 mM Tris-HCl pH 8, 250 mM NaCl, 8 M urea, 100 μM ZnCl_2_), lysed by sonication, clarified by centrifugation, and purified by Ni^2+^-affinity chromatography with GB1-TAZ2 refolding occurring on the column by washing with lysis buffer lacking urea (IMAC Sepharose, Cytiva). The eluent was reduced with the addition of β-mercaptoethanol (βME) to 50 mM, diluted two-fold with lysis buffer containing 1 mM ZnCl_2_, and then incubated with 200 U thrombin overnight at 4°C. Cleaved TAZ2 samples were diluted three-fold with a low-salt buffer (20 mM Tris-HCl pH 8, 5 mM βME, 10 μM ZnCl_2_), and purified by ion exchange chromatography (SP Sepharose, Cytiva).

For purification of GB1-MITF and GB1-E1A_CR1_, cell pellets were resuspended in native lysis buffer (20 mM Tris-HCl pH 8, 250 mM NaCl, 5 mM βME) and purified by Ni^2+^-affinity chromatography. The eluent was dialyzed overnight at 4°C (20 mM Tris pH 8, 5 mM βME), and if required 150 µg TEV protease was added to remove the affinity tag. Cleaved MITF and E1ACR1 were separated from His_6_-GB1 by Ni^2+^ affinity and ion exchange chromatography (Q Sepharose; Cytiva). All protein purifications were monitored by SDS-PAGE.

### Pulldown assay

GB1 and GB1-MITF fusion proteins (20 nmoles) were immobilized onto 20 μL IgG agarose beads (Cytiva) in pulldown buffer (20 mM Tris-HCl pH 8, 25 mM NaCl, 5 mM βME, and 10 μM ZnCl_2_). After washing away excess protein the beads were incubated with 20 nmoles of TAZ2 for 30 min. Following incubation, the beads were washed three times with pulldown buffer with or without the presence of 20 nmoles E1A_CR1_. After washing, IgG agarose beads were resuspended in Laemmli buffer and analyzed by SDS-PAGE.

### Isothermal titration calorimetry

Experiments were performed at 30 °C in 20 mM mM MES pH 6, 50 mM NaCl, 5 mM TCEP, and 10 μM ZnCl_2,_ using a VP-ITC microcalorimeter (MicroCal). TAZ1 or TAZ2 at a concentration of 800 μM was loaded into the syringe and injected into a calorimetric reaction cell containing 80 μM of E1A_CR1_, MITF_81-204_ or MITF_81-204 ΔDDVIDDII_. ITC experiments were collected in duplicate with 30 injections of 10 μL increments at 300 sec equilibration intervals. Thermograms were fit to a one-site binding model using the MicroCal Origin 7.0 Software.

### NMR spectroscopy

NMR spectra of MITF_81-204_ (400 µM) were collected in 20 mM 20 mM MES pH 6.0, 50 mM NaCl, and 5 mM DTT at 25 °C using a Varian INOVA 600 MHz spectrometer with a room temperature probe (Queen’s University, Kingston ON). Resonance assignments of MITF_81-204_ were determined by interpreting ^1^H-^15^N HSQC, HNCACB, CBCACONH, H(CCO)NH, (H)C(CO)NH, HNCO, and HN(CA)CO experiments. To assess binding between MITF and domains of CBP/p300, 10 µM ZnCl_2_ was added to the sample buffer and ^1^H-^15^N HSQC spectra were collected of 100 µM ^15^N-labelled MITF_81-204_ or MITF_110-161_ in the absence and presence of 200 µM TAZ1, or TAZ2.

All other NMR samples were prepared in MES pH 6.0, 5 mM βME, 10 μM ZnCl_2_, and 5% D_2_O and data was collected at 35 °C. For competition experiments, samples of 100 µM ^15^N-labelled MITF_110-161_ were prepared with or without 100 µM TAZ2 and 150 µM E1A. NMR spectra were acquired on a Bruker Avance III 700 MHz spectrometer equipped with a cryogenically cooled probe at the National Research Council Institute for Marine Biosciences (NRC-IMB, Halifax, NS). To determine the structure of the MITF_TAD_:TAZ2 complex, samples were prepared containing either 1 mM uniformly ^13^C/^15^N-labelled MITFT_TAD_ and 1.2 mM TAZ2, or 950 µM uniformly ^13^C/^15^N-labelled TAZ2 and 1150 µM MITF_TAD_. Resonance assignments were determined using standard triple resonance experiments and distance restraints for the MITF_TAD_:TAZ2 complex and distance restraints were obtained from ^15^N HSQC-NOESY, aliphatic and aromatic ^13^C HSQC-NOESY, and ^12^C/^14^N-filtered ^13^C-edited NOESY experiments. All NOESY experiments were acquired with a 100 ms mixing time.

Raw data were processed using NMRPipe (28) and analyzed using CcpNmr Analysis (29). After resonance assignment, secondary structure propensity (30) was calculated and chemical shift changes (Δ δ) upon addition of MITF_TAD,_ TAZ1 or TAZ2 were quantified as Δ δ = [(0.17Δδ_N_)^2^+ (Δδ_HN_)^2^]^1/2^ (31). For structure calculation NOESY peak lists and DANGLE-generated dihedral angle predictions (32) were provided to ARIA2 (33). After the initial fold of the MITF_TAD_:TAZ2 complex became apparent, zinc coordination restraints were incorporated for known zinc-binding residues of TAZ2 (34) and hydrogen bond distance restraints (1.8 ≤ *d*_OH_ ≤ 2.2 Å; 2.7 ≤ *d*_ON_ ≤ 3.2 Å) were applied to helical regions of the complex. In total 100 structures were generated over 8 iterations of automated NOE assignment, with the 20 lowest energy structures being selected for automated water refinement. The quality of the final ensemble of structures was assessed using the protein structure validation suite (35), and the Protein Data Bank validation tools. The MITF_TAD_:TAZ2 ensemble was deposited into the Protein Data Bank (accession no. 8E1D), while the chemical shift assignments of MITF81-204 and the MITFTAD:TAZ2 complex were deposited into the BMRB (accession nos. 51550 and 31038, respectively). PyMOL (Schrödinger Inc.) was used to interpret structures and generate figures.

### Peptide array

Peptide array experiments were performed as described previously (36). Briefly, MITF peptides were generated via automatic SPOT synthesis (37, 38). Peptides were synthesised on continuous cellulose membrane supports using 9-fluorenylmethyloxycarbonyl chemistry (Fmoc) by the MultiPep RSi Robot (Intavis). Arrays were pre-activated in absolute ethanol, followed by blocking in 5% BSA in 1x Tris-buffered saline with 0.1% tween-20 (TBS-T) for 4 hours at room temperature. GB1-TAZ2 was diluted to 2 µM using 200 mM NaCl, 50 mM Tris Base, 5% glycerol, pH 7.4, and overlaid onto the MITF array overnight at 4 °C. GB1-TAZ2 binding to MITF peptides was determined utilising a polyhistidine monoclonal mouse antibody (1:5000, Sigma, H1029), incubated on the array for 4 hours at 4 °C. Following this, a mouse horse radish peroxidase conjugated secondary antibody (1:5000) was incubated for 1 hour at room temperature. Antibodies were diluted into TBS-T with 1% BSA. Finally, binding signal was determined by enhanced chemiluminescence detection. All protein and antibody incubation steps were carried out under gentle agitation, and arrays washed following protein and antibody incubation steps 3 times in TBS-T.

### Luciferase-based transactivation assays

HEK 293A cells were cultured at 37°C with 5% CO_2_ in Dulbecco’s modified Eagle’s medium supplemented with 10% fetal bovine serum. For reporter assays, cells were seeded into 24 -well plates and transfected the following day at 70-80% confluency using jetPRIME transfection reagent. For each luciferase assay, a total of 500 ng of plasmid per well was transfected including 350 ng luciferase reporter (p5xGAL4-luc), 50 ng internal control (pCMV-Renilla), and 100 ng of expression plasmid (pGAL4-MITF). For co-transfections of involving MITF, 50 ng pGAL4-MITF_81-204_ and up 50 ng of pcDNA-p300 or pGFP-E1A_CR1_ were used. Luciferase activity was determined 18-24 h post-transfection using the dual-luciferase reporter assay system according to the manufacturers protocols. All luciferase values are an average of *Renilla* normalized luminescence and represent at least three independent transfections. Statistical significance was measured using one-way ANOVA followed by Dunnett’s multiple comparison test with a significance threshold *p*-value ≤ 0.05 and variation reported as standard error.

## RESULTS AND DISCUSSION

### MITF interacts with the TAZ1 and TAZ2 domains of CBP/p300

A combination of luciferase and pulldown experiments were used to investigate the MITF transactivation domain first characterized by Sato *et al*. (1997). Luciferase-based mammalian one-hybrid assays with MITF regions fused to the GAL4 DNA-binding domain demonstrated that transfection of plasmids containing the N-terminus of MITF, particularly pGAL4-MITF_1-204_ and pGAL4-MITF_81-204_, activated transcription 100-fold and 135-fold, respectively, compared to expression of pGAL4 alone (Figure 2A). In contrast pGAL4-MITF_1-100_ and pGAL4-MITF_289-419_ were unable to activate transcription above levels observed for pGAL4. These results are consistent with MITF containing a transactivation domain in its N-terminal sequence (17), and we do not observe evidence for transactivation from C-terminal fragments of MITF. Since transcriptional activation by MITF is partly mediated by CBP/p300 recruitment (39), we investigated the coactivation potential of p300 by transfecting pGAL4-MITF_81-204_ alone or with increasing amounts of pcDNA-p300. We found co-transfection of pcDNA-p300 potentiated transactivation by pGAL4-MITF_81-204_ up to 5-fold, indicating that we are observing a functional interaction between MITF and CBP/p300 (Figure 2B).

**Figure 2.**
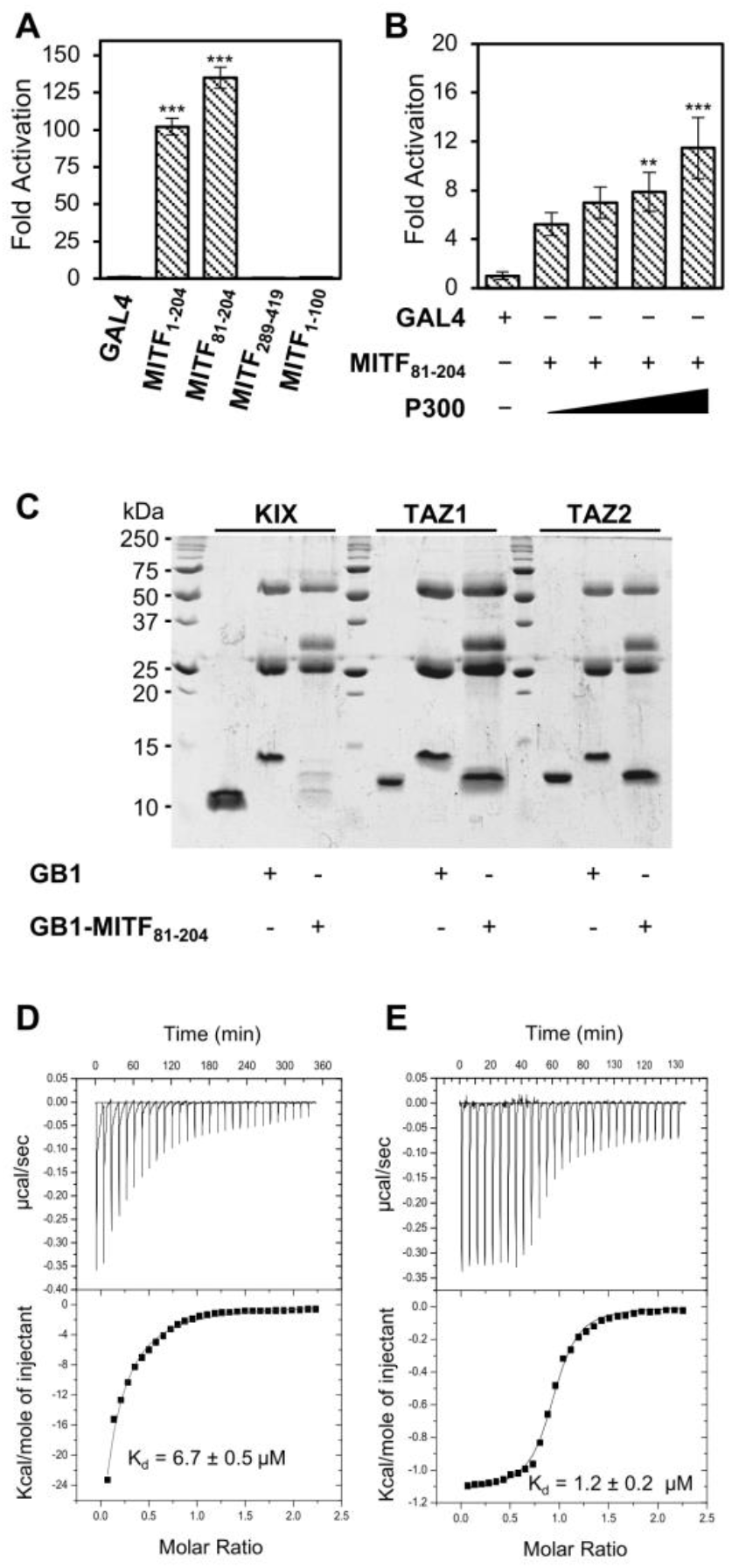
MITF interacts with CBP/p300 through TAZ1 and TAZ2. **(A)** Luciferase-based mammalian one-hybrid transactivation assays performed in HEK 293A cells using 100 ng of indicated pGAL4-MITF fusion proteins. **(B)** Luciferase-based mammalian one-hybrid transactivation assays performed for 50 ng of pGAL4-MITF_81-204_ with co-transfection of up to 50 ng of p300. Asterisks indicate statistical significance by one-way ANOVA and Dunnetts multiple comparison test (** *p* ≤ 0.01, *** p ≤ 0.005) compared to pGAL4 and variation is reported as SEM.**(C)** 15% SDS-PAGE analysis of the ability of immobilized GB1 or GB1-MITF_81-204_ to pulldown and interact with purified recombinant KIX, TAZ1, and TAZ2. ITC thermograms of 800 µM TAZ1 **(D)** or TAZ2 **(E)** titrated into 80 µM MITF_81-204_ and fit to a one-site binding model with measured dissociation constants (K_d_) indicated.

CBP/p300 is a large protein with multiple protein-interaction domains, and outside of the histone acetyltransferase region, the TAZ1, TAZ2, and KIX motifs are candidate interaction sites for a range of intrinsically disordered activation domains of cellular transcription factors including STAT1, STAT2, p53 and *c-Myb* (40–43). To corroborate our transactivation assays and to determine the regions of CBP/p300 that interact with MITF, we completed pulldown assays using various fragments of MITF expressed as fusion proteins with the B1 binding-domain of *Streptococcus* protein G (GB1). GB1-MITF fusion proteins were expressed in *E. coli*, purified, and then bound to IgG Sepharose beads and mixed with an equivalent amount of purified recombinant KIX, TAZ1, or TAZ2. SDS-PAGE analysis of bound protein indicates that GB1-MITF_81-204_ interacts with both the TAZ1 and TAZ2 domains of CBP/p300 (Figure 2C). Using isothermal titration calorimetry, we measured dissociation constants (K_d_)’s of 6.7 ± 0.5 µM and 1.24 ± 0.23 µM for the association of MITF_81-204_ with TAZ1 and TAZ2, respectively (Figure 2D,E). The association of TAZ2 with CBP/p300 is consistent with pulldown studies between larger fragments of CBP/p300 and MITF (17); however, to our knowledge the association between TAZ1 and MITF has not been previously reported.

### MITF interacts with TAZ1 and TAZ2 via a common transactivation domain

Since MITF_81-204_ demonstrated both a functional interaction with p300 in HEK 293A cells and direct binding to TAZ1 and TAZ2 by pulldown assay, nuclear magnetic resonance (NMR) spectroscopy was used to further characterize the molecular properties of MITF and its interaction with these domains. The ^1^H-^15^N HSQC spectrum of ^15^N-labelled MITF_81-204_ has the expected number of peaks with sharp linewidths and little chemical shift dispersion (Figure S1A), which is indicative of MITF being an intrinsically disordered protein (44). Standard triple resonance experiments were used to assign 122/124 residues and 96% of the backbone resonances of MITF_81-204_. Secondary structure propensity analysis indicates that although disordered, MITF_81-204_ has some propensity to form an α-helix, with residues Met105-Gln115, Ile124-Met144, Gly152-Gly159, and Cys188-Ala198 having the highest SSP scores (Figure S1B).

We then used NMR-based chemical shift perturbation studies to clearly define which residues within MITF_81-204_ bind TAZ1 and TAZ2 (Figures 3A and S2). When compared to free peptide, most resonances of MITF_81-204_ experienced minor chemical shift perturbations upon the addition of unlabelled TAZ1 or TAZ2 (Figures 3B and S2), with no obvious change in linewidth or peak intensity. However, residues Gln115-Ala141 of MITF undergo line broadening to the extent that peaks disappear from the ^1^H-^15^N HSQC spectrum upon addition of TAZ2, with similar changes observed for TAZ1 (Figure S2), which is indicative of an intermolecular interaction occurring in this region.

**Figure 3.**
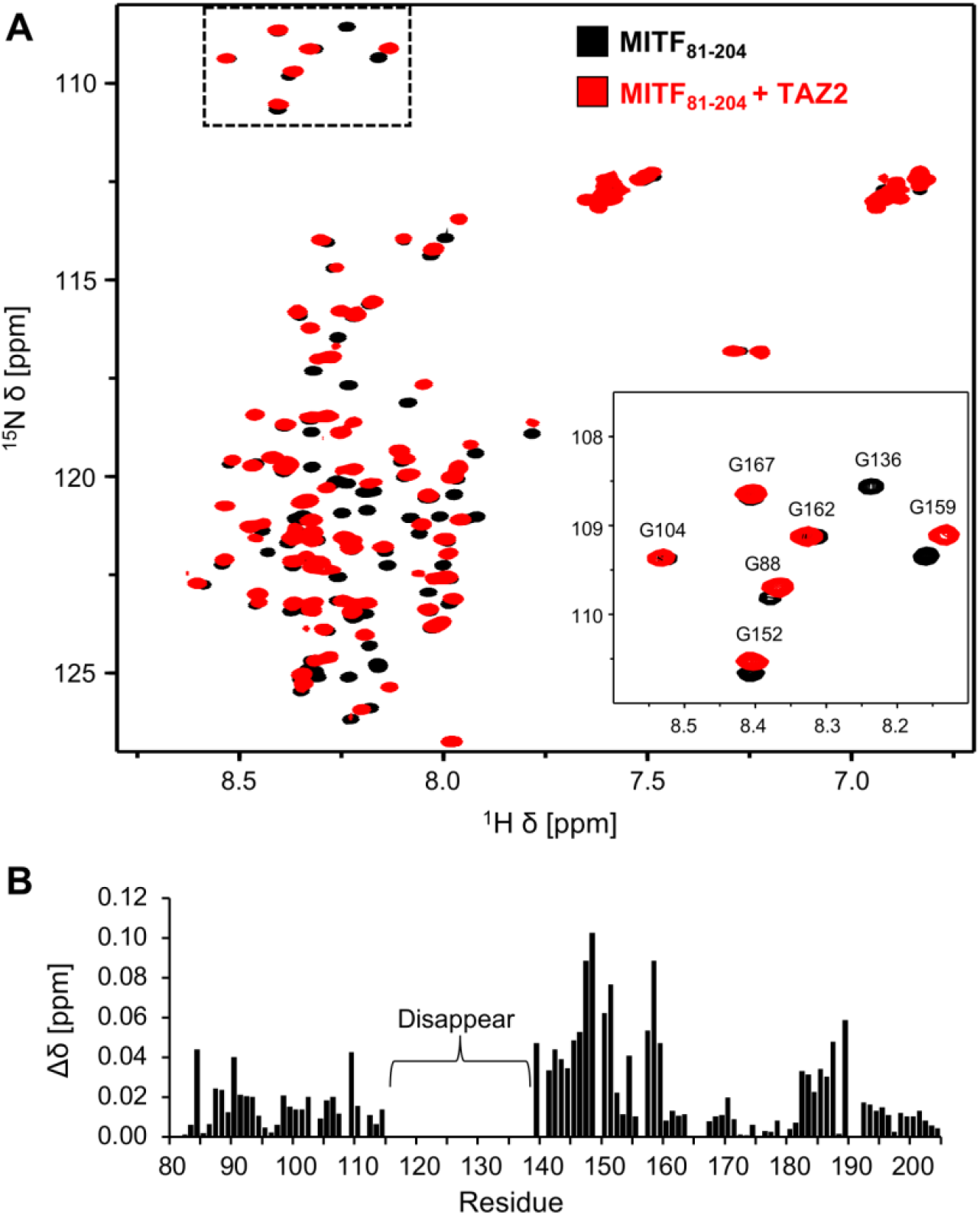
MITF binds TAZ2 through MITF_TAD_ **(A)** ^1^H-^15^N HSQC of 100 µM ^15^N-labelled MITF_81-204_ overlayed in the absence (black) and presence (red) of 200 µM unlabelled TAZ2. Residue assignments of a subset of peaks are indicated in the insert. **(B)** The chemical shift changes (Δ δ = [(0.17Δδ_N_)^2^+ (Δδ_HN_)^2^]^1/2^) that each residue of MITF_81-204_ experienced upon addition of TAZ2 results are plotted, with peaks that disappear upon addition of TAZ2 indicated.

To improve spectral quality and reduce resonance overlap, we titrated ^15^N-labelled MITF_110-161_ with TAZ1 and TAZ2. When free in solution MITF_110-161_ has low peak dispersion and sharp resonances, suggesting that it is disordered (Figure S3A). With this smaller construct more MITF resonances remained visible upon the addition of TAZ1 or TAZ2, with the addition of TAZ2 often resulting in larger chemical shift changes than for TAZ1 (Figure S3). Interestingly, resonances of MITF experience similar magnitude and direction of chemical shift changes upon the addition of either TAZ1 or TAZ2, suggesting that MITF adopts a similar structure when binding these proteins. Consistent with the peak disappearance observed for MITF_81-204_, residues Arg110-Ala141 of MITF (hereafter referred to as MITF_TAD_) underwent the most significant chemical shift changes, while the rest of the peptide experienced only minor chemical shift changes. All resonances of MITF_110-161_ are observable when bound to TAZ2 but not when bound to TAZ1 (Figure S3), as such we used NMR spectroscopy to characterize the structure of the MITFTAD:TAZ2 complex.

### Structure of the MITF_TAD_:TAZ2 complex

To determine the structure of the MITF_TAD_:TAZ2 complex, resonance assignments were determined for ^13^C/^15^N-labelled TAZ2 in complex with MITF_TAD_, as well as ^13^C^15^N-labelled MITF_TAD_ saturated with TAZ2 (Figure 4A). These spectra were high quality and allowed for assignment of 97%, 81%, and 56% of backbone, sidechain, and aromatic resonances, respectively. When bound to TAZ2, residues Asp118-Asp139 of MITF have ^15^N{^1^H} NOE values > 0.25, with most of these values ranging from 0.5 to 0.6 (Figure 4B). Many of these residues (Asp121, Asp122, and Ser125-Ser129) have chemical shift index (Figure 4C) values that suggest this region forms an α-helix upon binding TAZ2.

**Figure 4.**
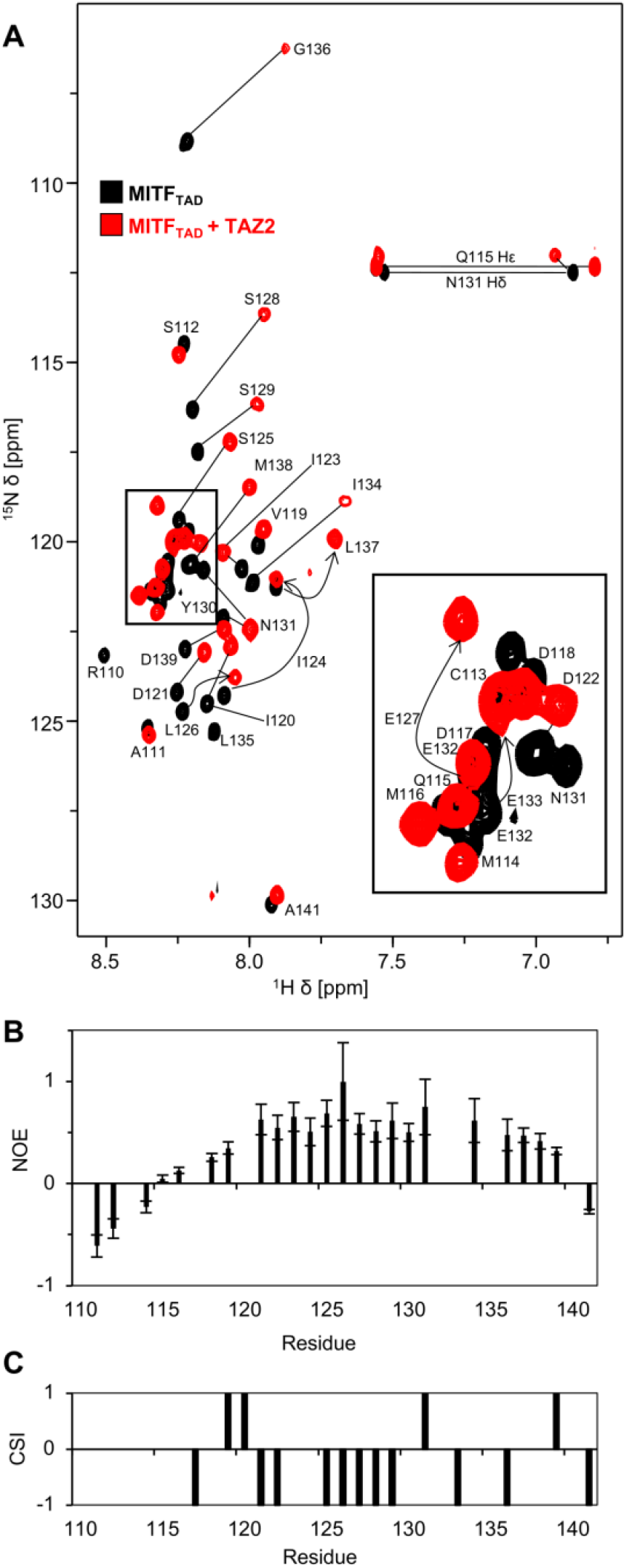
Characterization of structure and dynamics of MITF_TAD_ bound to TAZ2 **(A)** Overlay of ^1^H-^15^N HSQC of 1 mM ^13^C/^15^N-labelled MITF_TAD_ in the absence (black) and presence (red) of 1.2 mM unlablled TAZ2 (red). Residue assignments of resonance peaks are indicated, and arrows indicate resonance shifts upon TAZ2 addition. **(B)** Plot of ^15^N{^1^H} NOE values for reach residue of MITF_TAD_ when bound to TAZ2. **(C)** Secondary structure propensity (SSP) values per residue of MITF_TAD_ when bound to TAZ2. Positive values indicate α-helical and negative values indicate β-sheet propensity.

After resonance assignment of MITF_TAD_:TAZ2, 1783 NOE-derived distance restraints were obtained from ^13^C-NOESY-HSQC, ^15^N-NOESY-HSQC, and ^12^C/^14^N-filtered ^13^C-edited NOESY spectra and with dihedral angle restraints were used to generate a 20-member structural ensemble of the MITF_TAD_:TAZ2 complex (Table 1 and Figure 5A). The resulting ensemble was high quality with 91.1%, 6.8%, and 2.1% of ordered residues having favoured, allowed, and disallowed dihedral angles. In this ensemble TAZ2 is well-converged while the structure of MITF is less defined. Analysis using protein data bank validation reports indicate that nearly all TAZ2 and a small region of MITF (residues Pro1727-Gln1811 and Leu126-Ile134, respectively) can be considered the ordered core of this complex, with a root-mean-square deviation (RMSD) of 0.7 Å and 1.1 Å for backbone and heavy atoms, respectively. This positioning of MITF_TAD_ is supported by observed intermolecular NOE contacts between Leu126-Ala1787, Tyr130-Ile1735/Leu1788, and Ile134-Pro1780/Ile1781/Gln1784 (Figure 5B). The structure of TAZ2 is consistent with previously determined structures of TAZ2 in isolation or in complex with transcription factors and consists of four alpha helices (α1-α4) spanning residues Gly1728-Gln1747, Cys1758-Thr1768, Pro1780-Ala1793, and Pro1804-Ile1809, respectively. MITF_TAD_ has two α-helices spanning residues Asp118-Ser129 Glu132-Leu135 that are supported by both cross peaks in NOESY spectra and the calculated CSI for MITF_TAD_. Although the presence of these α-helices is well-defined there are few intermolecular contacts with TAZ2 in these regions, making the positioning of these α-helices with respect to TAZ2 somewhat variable. MITF_TAD_ binds a hydrophobic surface of TAZ2 created by the intersection of α1-α3 (Figure 5C), which is also targeted by other transcription factors, such as p53 and viral E1A (45, 46). The ^15^N{^1^H} NOE values of MITF_TAD_ in complex with TAZ2 are lower than for a well-ordered protein. This is consistent with the relatively few intermolecular NOESY contacts that were observed and suggests that the MITF_TAD_:TAZ2 complex may be somewhat dynamic and lacks a single low-energy bound state. Intrinsically disordered proteins such as MITF often form a dynamic or fuzzy complex when interacting with binding partners, and recently this has been observed for the E7 protein binding to TAZ2 (47).

**Table 1.** Statistics of the NMR-derived ensemble of MITF_TAD_:TAZ2.

|  |  |
| --- | --- |
| **Completeness of resonance assignments (%)<sup>a</sup>** |  |
| Backbone | 97 |
| Side Chain | 81 |
| Aromatic | 56 |
| **Number of conformational restraints** |  |
| Total NOE restraints | 1783 |
| Intra-residue (i = j) | 922 |
| Sequential (\|i - j\| = 1) | 402 |
| Medium range (1 < \|i - j\| < 5) | 249 |
| Long range (\|i - j\| ≥ 5) | 154 |
| Intermolecular | 15 |
| Ambiguous | 41 |
| Dihedral angle restraints | 190 |
| Hydrogen bond restraints | 90 |
| NOE restraints per residue | 13.1 |
| Long range restraints per residue | 1.1 |
| **Residual restraint violations<sup>b</sup>** |  |
| RMSD NOE restraints (Å) | 0.05 |
| RMSD dihedral angle restraints (°) | 0.34 |
| Mean distance violations per structure |  |
| > 0.3 Å | 3.6 |
| > 0.5 Å | 0.15 |
| Mean dihedral angle violations per structure > 5° | 0.05 |
| **Model quality<sup>c</sup>** |  |
| RMSD backbone atoms (Å)<sup>d</sup> | 0.7 |
| RMSD heavy atoms (Å)<sup>d</sup> | 1.1 |
| RMSD bond lengths to ideal geometry (Å) | 0.006 |
| RMSD bond angles to ideal geometry (°) | 0.8 |
| **Ramachandran plot statistics<sup>de</sup>** |  |
| most favored regions (%) | 91.1 |
| allowed regions (%) | 6.8 |
| disallowed regions (%) | 2.1 |
| **Global quality scores (Raw / Z-score)<sup>c</sup>** |  |
| Verify3D | 0.07 / -6.26 |
| ProsaII | 0.64 / -0.04 |
| ProCheck G-factor (phi-psi)<sup>d</sup> | -0.15 / -0.28 |
| ProCheck G-factor (all)<sup>d</sup> | -0.39 / -2.31 |
| MolProbity clash score | 45.41 / -6.27 |
| **Model contents** |  |
| Ordered residue ranges<sup>f</sup> | 1727-1811 |
|  | 126-134 |
| Total no. of residues | 136 |
| BMRB accession no. | 31038 |
| PDB accession no. | 8E1D |

<sup>a</sup>Calculated using RCSB validation server
<sup>b</sup>Calculated using CNS version 1.21
<sup>c</sup>Calculated using PSVS version 1.5
<sup>d</sup>Calculated over ordered residue ranges
<sup>e</sup>Calculated using Molprobity through PSVS version 1.5
<sup>f</sup>Based on core residue ranges listed in RCSB validation report.

**Figure 5.**
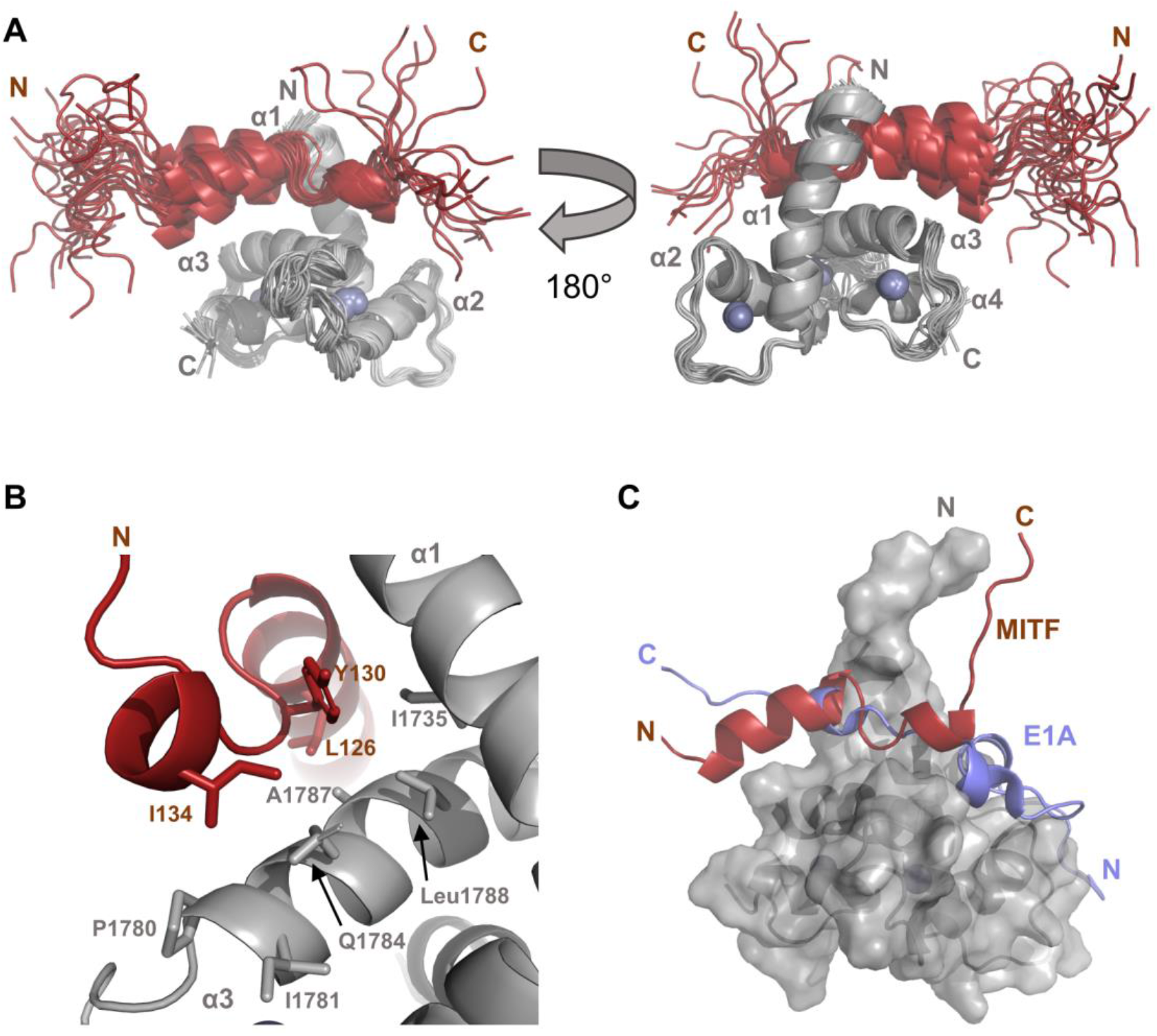
Structure of MITF_TAD_:TAZ2 complex. **(A)** Backbone ribbon illustration of the 20-member structural ensemble of the MITF_TAD_:TAZ2 complex, with TAZ2 coloured grey and MITF_TAD_ shown in red. The disordered N-terminal residues of TAZ2 are omitted for clarity and zinc ions are shown as spheres. **(B)** Close-up view of the interaction interface between MITFTAD and TAZ2. Residues with observable intermolecular NOE contacts are shown as sticks and labelled. **(C)** Superposition of E1A (PDB accession no. 2KJE) with the MITF_TAD_:TAZ2 complex. TAZ2 is shown as a transparent grey surface and E1A is coloured blue.

### An acidic motif mediates binding of MITF_TAD_ to TAZ2

To identify residues important for the MITF_TAD_:TAZ2 interaction in a high-throughput manner, a peptide array was used to probe for variation in the ability of MITF to pull down TAZ2. Due to technical limitations a shorter MITF peptide, MITF_117-135_, was used for this assay, which corresponds to the region of MITF that associates directly with TAZ2 based on the NMR-derived structure and heteronuclear NOE measurements. Each residue of MITF_117-135_ was sequentially mutated to alanine, probed with GB1-TAZ2, and visualized using chemiluminescence. Although no point mutations completely ablated GB1-TAZ2 binding, decreased interaction was observed when residues within Asp117-Asp122 (DDVIDD) were mutated to alanine. This overlaps with the ΦXXΦΦ motif that is conserved in MiT/TFE members, suggesting that this motif is necessary for association with TAZ2 (Figure 6A).

**Figure 6.**
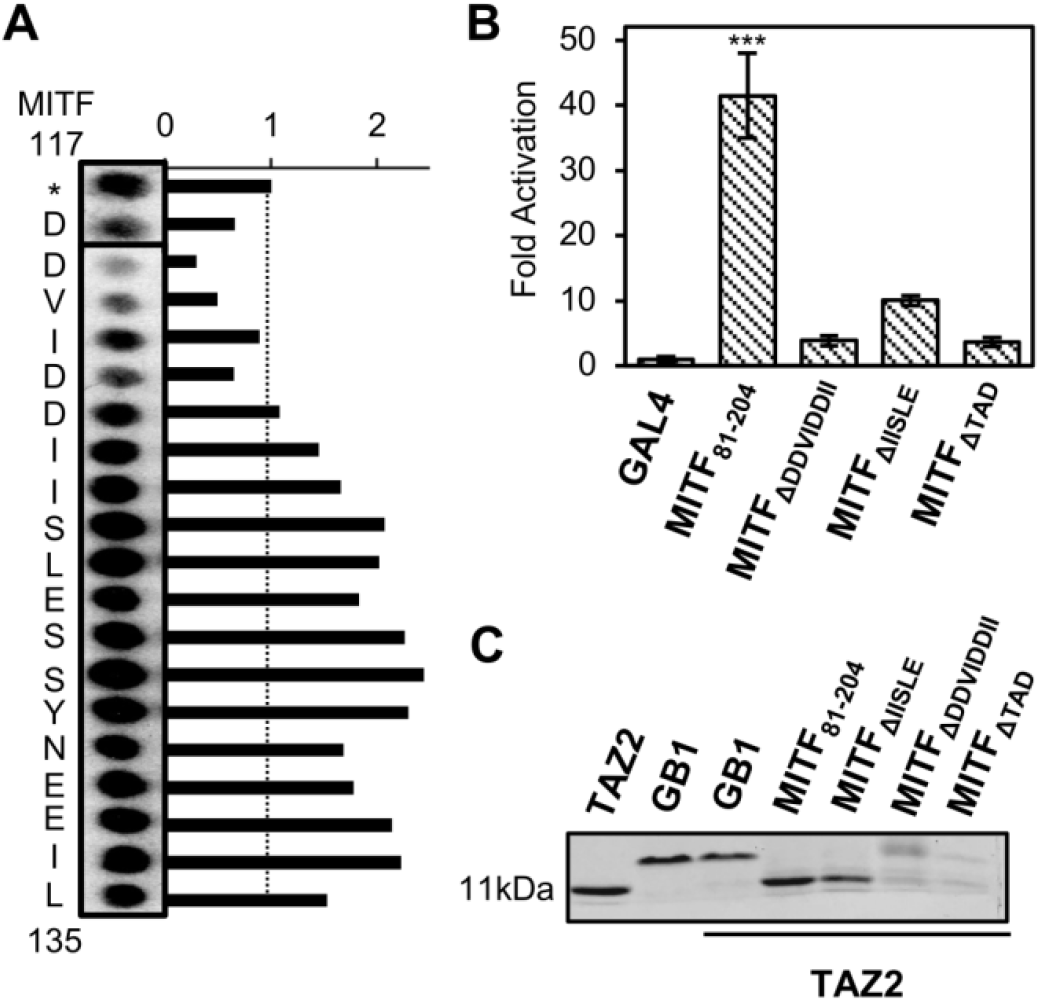
MITF_TAD_ mutagenesis ablates TAZ2 binding. **(A)** Peptide microarray of MITF_117-135_ with each residue sequentially mutated to alanine, probed for its binding affinity with GB1-TAZ2 and visualized by chemiluminescence. * Annotates wildtype MITF_117-141_ to which all peak intensities were normalized. **(B)** Luciferase-based transcriptional activity after transfection of cells with 100 ng pGAL4-MITF_81-204_ (either wildtype or containing the indicated deletion). Statistical significance by one-way ANOVA and Dunnetts multiple comparison test compared to Gal4 is indicated (*** p ≤ 0.005) and variation is reported as SEM. **(C)** Protein pulldown assay of GB1 and GB1-MITF_81-204_ analyzed by SDS-PAGE. Reference bands are shown for TAZ2 and GB1, while pulldown of TAZ2 by the indicated GB1-MITF_81-204_ deletion mutants is shown.

These peptide array results are consistent with luciferase-based mammalian-one hybrid assays and pulldown experiments (Figure 6B,C), where this acidic motif between Asp117-Ile124 (DDVIDDII) was found to be necessary for both transcriptional activation and pulldown of TAZ2 by MITF_81-204._ Supporting this, mutation of this motif rendered the interaction between TAZ2 and MITF_81-204_ undetectable by isothermal titration calorimetry (Figure S5). Conversely, mutation of individual residues between Ile123-Glu127 (IISLE) did not decrease the observed binding in the peptide array and complete removal of Ile123-Glu127 resulted in only partial disruption of TAZ2 pulldown and transactivation by MITF_81-204_. This is consistent with studies that show deletion of IISLE significantly reduces the function of the MITF activation domain, but retains the ability to activate transcription above the GAL4 reporter alone (17). An interesting finding is that even though Ile123-Glu127 (IISLE) are relatively well-ordered in the MITF_TAD_:TAZ2 complex, these residues are not primary modulators of the interaction. Instead, an acidic motif (DDVIDD) that remains somewhat dynamic when interacting with TAZ2 is a primary modulator of TAZ2 binding and transactivation potential. Hence, electrostatic interactions between MITF and TAZ2, which is basic, are important modulators of this interaction.

### Adenoviral peptide E1A competes with MITF for the same TAZ2 binding surface

Since MITF interacts with TAZ2 through a large interaction surface instead of a distinct binding pocket, the ability of a peptide to block the MITF:TAZ2 interaction was evaluated. The adenovirus early region 1 A (E1A) plays a critical role in viral infections by deregulating the hosts ’ cell cycle through interactions with essential cellular proteins including CBP/p300 (24). The interaction between CBP/p300 and E1A is mediated by the TAZ2 domain and a region within conserved region 1 of E1A (residues 54-82; E1A_CR1_) binds the same surface of TAZ2 as MITF_TAD_ (Fig 5C), suggesting that E1A, MITF, and other transcription factors may potentially compete for the TAZ2 domain of CBP/p300 to mediate viral infection (17, 39, 48, 49).

To test if E1A competes with MITF for TAZ2, an NMR-based displacement experiment was performed. Unlabelled TAZ2 was initially titrated into ^15^N-labelled MITF_110-161_ and HSQC spectra were collected. The presence of TAZ2 resulted in significant chemical shifts of many of the backbone ^1^H-^15^N resonances in the spectrum consistent with the formation of a TAZ2:MITF complex (Figure 7A). Upon the subsequent addition of unlabelled E1A_CR1_ to this sample, the chemical shifts of MITF_110-161_ became more similar to those of the unbound peptide, indicating that E1A_CR1_ can displace MITF_110-161_ from TAZ2. These findings are supported by pulldown experiments, where incorporating E1A_CR1_ into the washes was sufficient to abrogate the ability of MITF_81-204_ to pulldown TAZ2 (Figure 7B). This also reflects isothermal titration calorimetry data that shows E1A_CR1_ can bind TAZ2 with ∼6-fold higher affinity than MITF_81-204_ (Figure S6).

**Figure 7.**
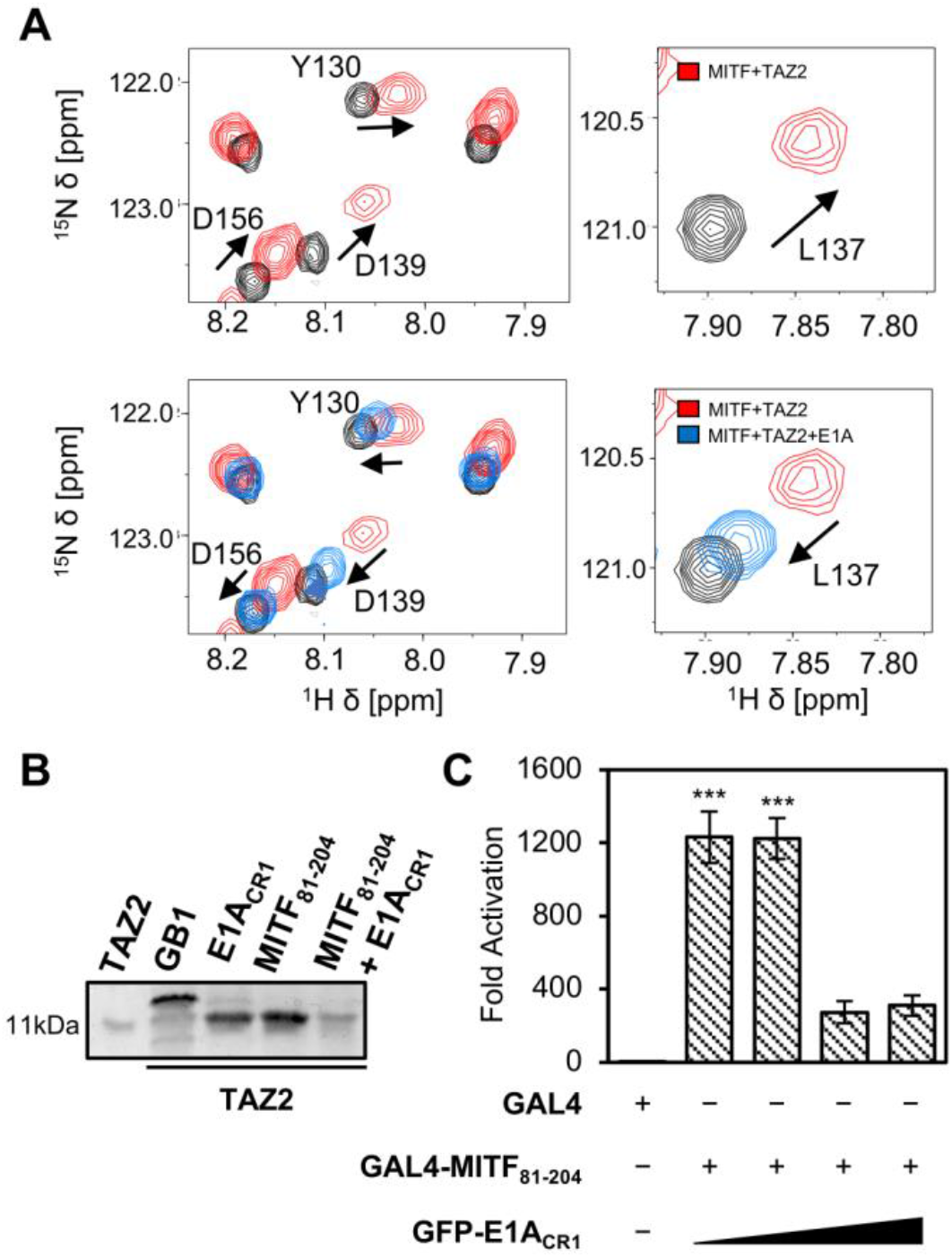
E1A_CR1_ competes with MITF for TAZ2. **(A)** Overlay of ^1^H-^15^N HSQC spectra of 100 µM ^15^N-labelled MITF_110-161_ (black), 100 µM MITF_110-161_ in the presence of 100 µM TAZ2 (red), and 100 µM MITF_110-161_ in the presence of both 100 µM TAZ2 and 150 µM E1A_CR1_ (blue). **(B)** TAZ2 pulldown by immobilized GB1, GB1-E1A_CR1_ and GB1-MITF_81-204_ as visualized using SDS-PAGE. Pulldown of TAZ2 by MITF_81-204_ when E1A_CR1_ is included in wash steps is shown in the righthand lane. **(C)** Luciferase-based transactivation assay of HEK 293A cells co-transfected with 50 ng pGAL4-MITF_81-204_ and up to 50 ng of pGFP-E1A_CR1_. Statistical significance by one-way ANOVA and Dunnetts multiple comparison test (*** *p* ≤ 0.005) is compared to activation from pGAL4, and variation is reported as SEM.

To assess whether MITF-dependent transcriptional activation could be suppressed by E1A_CR1_, we performed luciferase-based one-hybrid assays in which pGAL4-MITF_81-204_ was co-transfected with increasing amounts of pGFP-E1A_CR1_ in HEK 293A cells. The addition of E1A_CR1_ reduced luciferase activation by 5-fold, providing functional inhibition of MITF-dependent transcriptional activation (Figure 7C) and indicating peptides that bind tightly to the α1-α3 surface of TAZ2 can act as effective inhibitors of MITF function.

### Plasticity of transcriptional co-activator recruitment by MITF

CBP/p300 is recruited to promoters where it may simultaneously bind different transcription factors at nearby binding sites. Long disordered regions linking the multiple domains of CBP/p300 provide flexibility to interact with binding partners through several protein-interaction domains. Particularly the KIX, TAZ1, and TAZ2 domains facilitate activation domain binding with a broad range of affinity and specificities. For example, the tumor suppressor p53 activation domain binds promiscuously to CBP/p300 with affinity for KIX, TAZ1 and TAZ2 (50). We have found that the MITF activation domain also recruits CBP/p300 by binding to multiple domains, with a moderate binding affinity for both TAZ1 (K_d_ 6.7 ± 0.5 µM) and TAZ2 (K_d_ 1.24 ± 0.23 µM). Multivalent binding of CBP/p300 domains to MITF may promote cooperative interactions and could facilitate functional interactions between MITF and other transcriptional regulators that interact with TAZ1 or TAZ2. This promiscuous transcription factor binding by CBP/p300 provides a rationale for how the functional outcome of MITF overexpression/suppression depends in part on the cellular context and which transcription factors other than MITF are present (3).

The disordered nature of activation domains within transcriptional complexes allows for recognition of a variety of different sequences, where a small number of residues can mediate interactions with a high specificity. This study identifies a small acidic motif (DDVIDD) that overlaps with the ΦXXΦΦ motif within the MITF activation domain that facilitates TAZ2 binding and MITF transcription. Given the proximity of serine residues to this motif, and that phosphorylation occurs within the activation domain promoting CBP/p300 interactions (51), it is possible that post-translational modifications modulate MITF affinity for TAZ2, thereby modulating MITF activity. Binding to TAZ2 also brings MITF near the catalytic HAT domain, which may provide an additional method of regulation through post-translational mechanisms of acetylation, where CBP/p300 may directly regulate the affinity of MITF for DNA (52).

In summary, we have determined the MITF activation domain binds both the TAZ1 and TAZ2 domains of CBP/300 and interacts with TAZ2 via a hydrophobic surface formed by α1-α3, corresponding to the same TAZ2 binding site as adenoviral E1A. Our data shows the MITF_TAD_:TAZ2 complex maintains some conformational flexibility and is modulated primarily by an acidic motif that can be disrupted by peptides that bind TAZ2 with high affinity. This work provides a detailed characterization of how MITF binds to CBP/p300 and outlines how peptides that target TAZ2 can inhibit MITF function.

## DATA AVAILABILITY

The atomic coordinates and structure of the MITF_TAD_:TAZ2 complex have been deposited into the Protein Data Bank under accession number 8E1D. The chemical shifts of the MITF_TAD_:TAZ2 complex and MITF_81-204_ were deposited into the BioMagResBank under accession numbers 31038 and 51550, respectively.

## SUPPLEMENTARY DATA

Supplementary data are available at NAR online.

## ACKNOWLEDGEMENTS

We thank Dr. Stephen Bearne for providing access to an isothermal titration calorimeter and Mr. Ian Burton from the National Research Council Institute for Marine Biosciences (NRC-IMB) for assistance with NMR data acquisition.

