## Supplementary Data for "Structural basis of CBP/p300 recruitment by the microphthalmia-associated transcription factor"

**Contents**: Supplementary figures S1-S6.

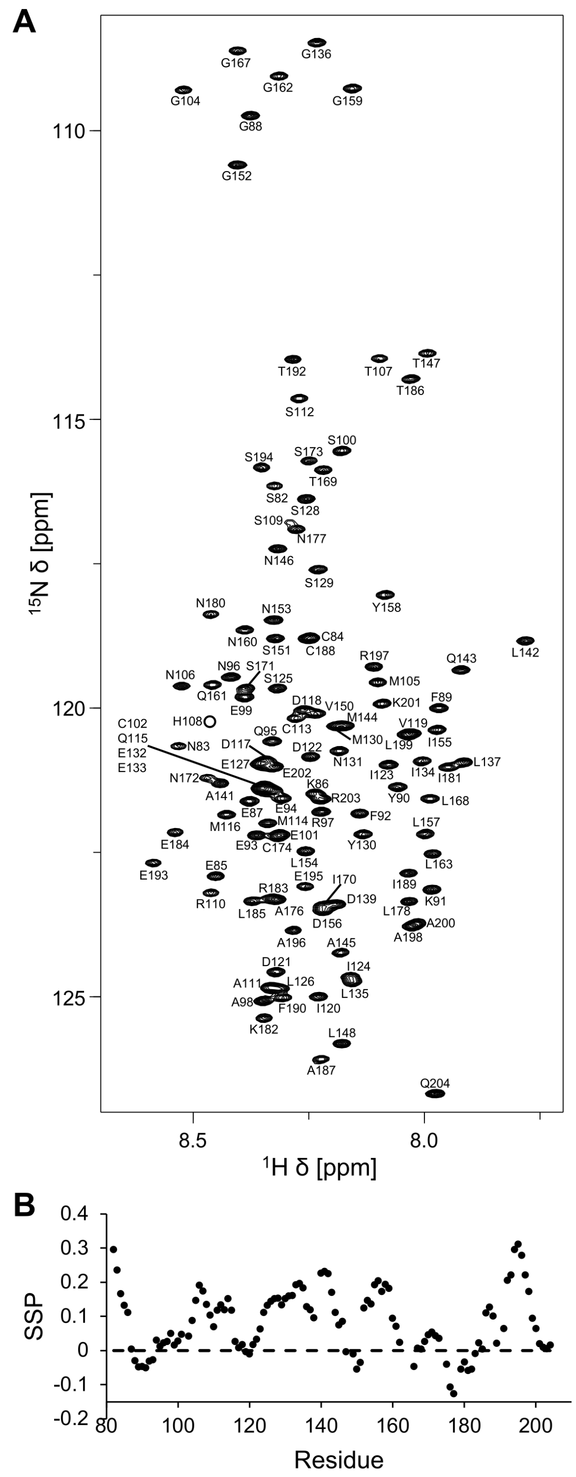


**Supplementary Figure S1. (A)** $^{1}$H-$^{15}$N HSQC of $^{13}$C/$^{15}$N-labelled MITF$_{81\text{-}204}$, residue assignments for backbone resonance peaks are indicated. **(B)** Secondary structure propensity (SSP) values per residue of MITF$_{81\text{-}204}$ calculated from Cα and Cβ chemical shifts.

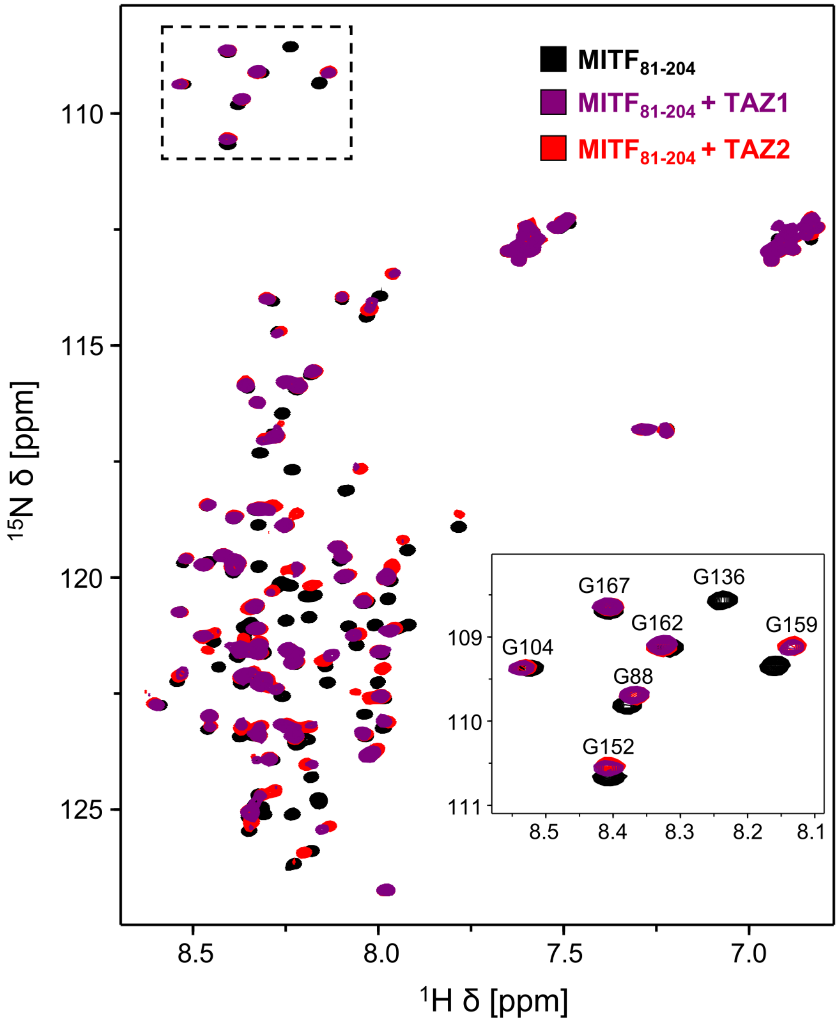


**Supplementary Figure S2.** $^{1}$H-$^{15}$N HSQC of 100 μM $^{15}$N-labelled $MITF_{81\text{-}204}$ in the absence (black) and presence of 200 μM of unlabelled TAZ1 (purple) or TAZ2 (red). Resonance assignments for a subset of peaks are indicated in the inset.

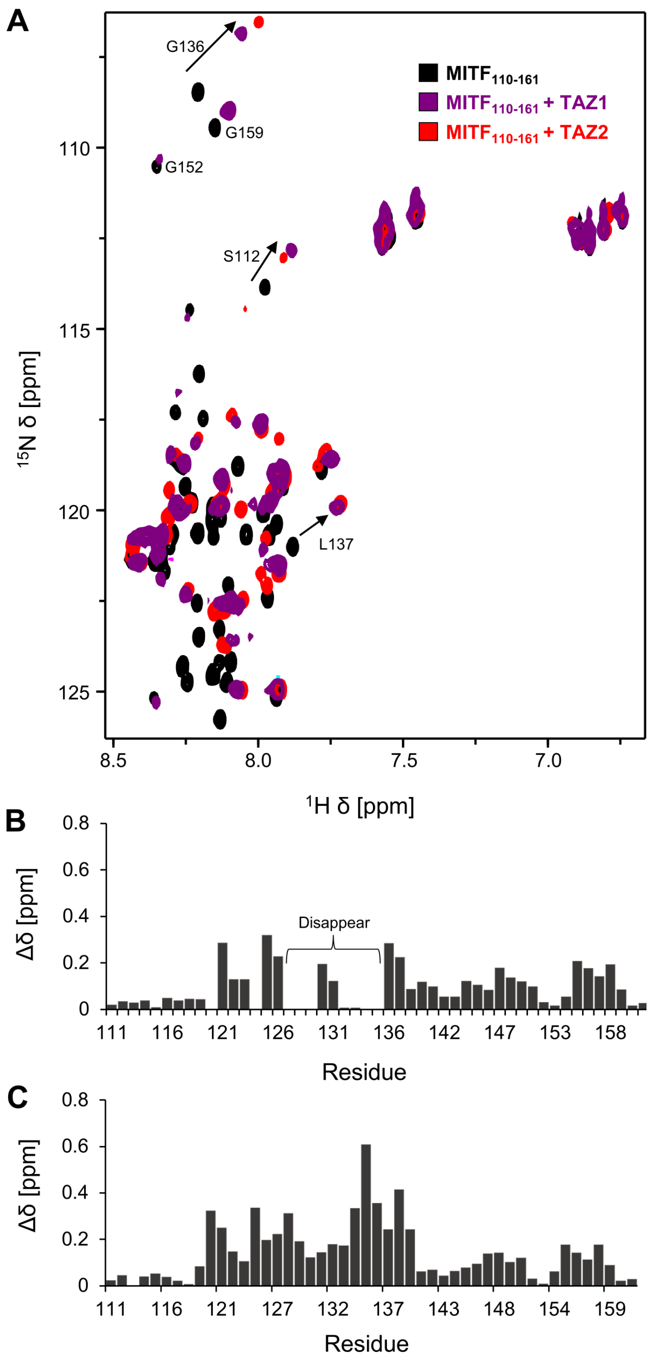


**Supplementary Figure S3. (A)** $^{1}$H-$^{15}$N HSQC of 100 μM $^{15}$N-labelled MITF$_{110\text{-}161}$ overlayed in the absence (black) and presence of 200 μM unlabelled TAZ1 (purple) or TAZ2 (red). Resonance assignments are indicated for non-overlapped peaks that shift upon addition of TAZ1 or TAZ2. Plots of chemical shift changes ($\Delta\delta = [(0.17\Delta\delta_N)^2 + (\Delta\delta_{HN})^2]^{1/2}$) that each residue of MITF$_{110\text{-}161}$ experienced upon addition of **(B)** TAZ1 or **(C)** TAZ2. Regions of MITF where resonances disappear upon addition of TAZ1 are indicated.

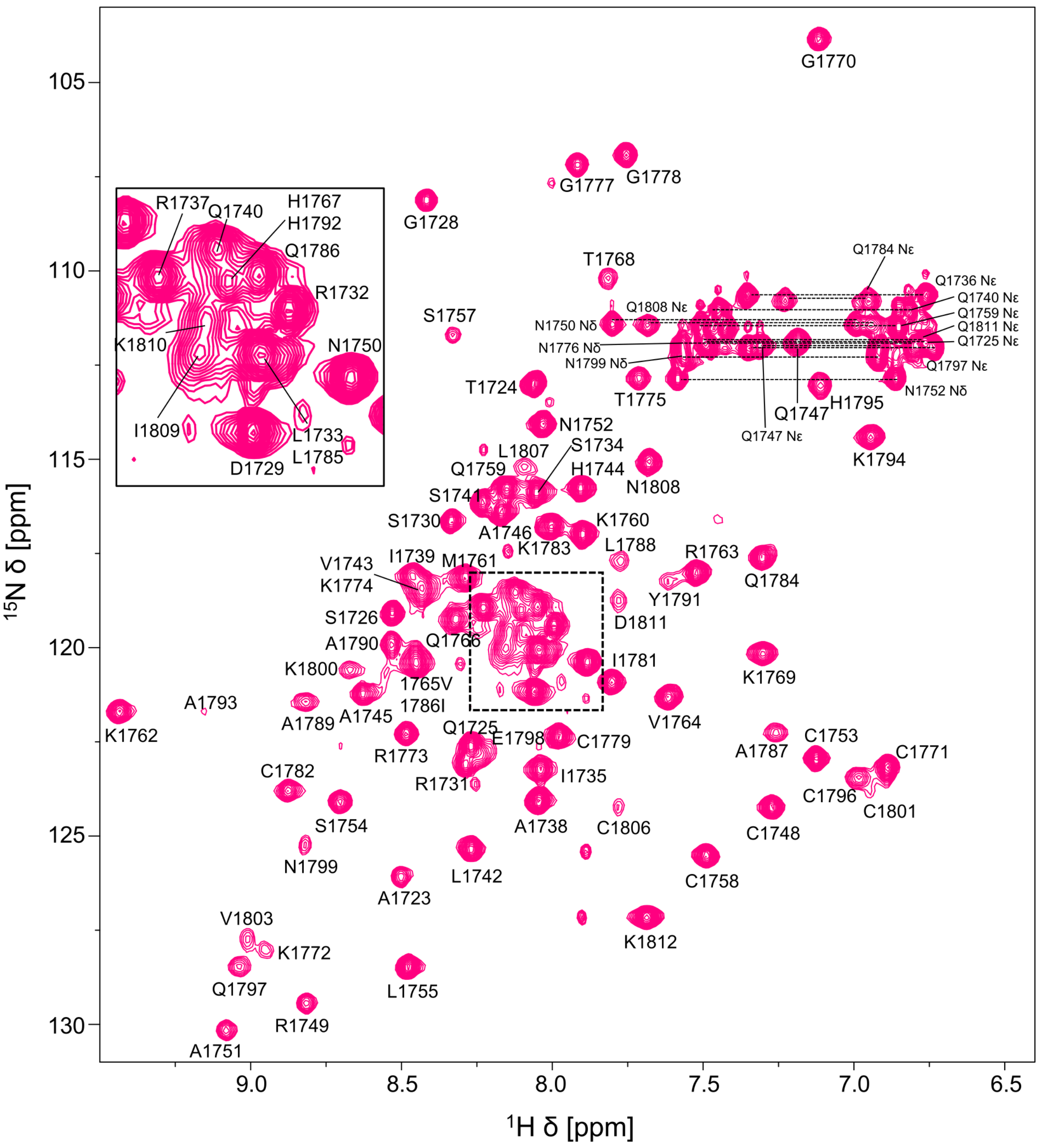


**Supplementary Figure S4.** $^1$H-$^{15}$N HSQC of 950 μM $^{13}$C/$^{15}$N-labelled TAZ2 in the presence of 1150 μM unlabelled MITF$_{TAD}$. Resonance assignments are indicated.

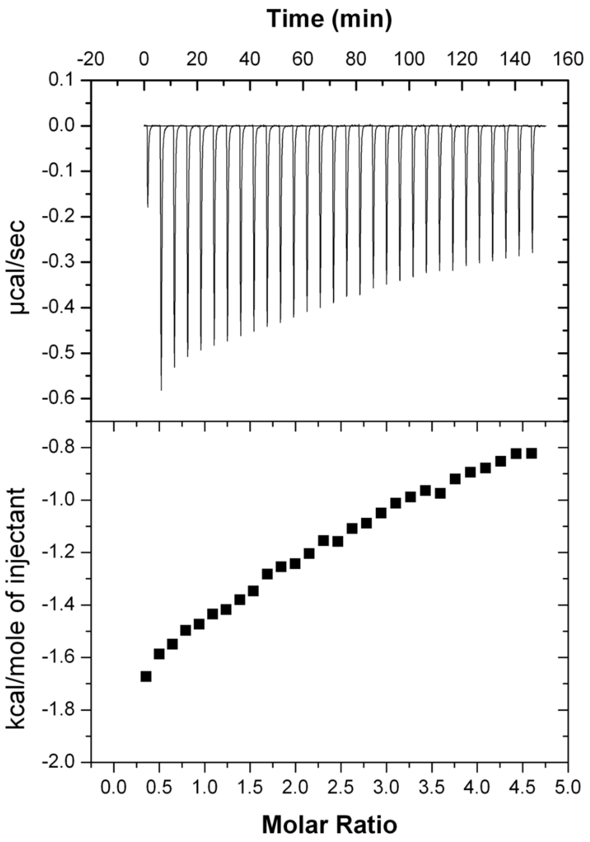


**Supplementary Figure S5.** Isothermal titration calorimetry thermogram of 800 µM TAZ2 titrated into 80 µM $MITF_{81\text{-}204\ \Delta DDVIDDII}$.

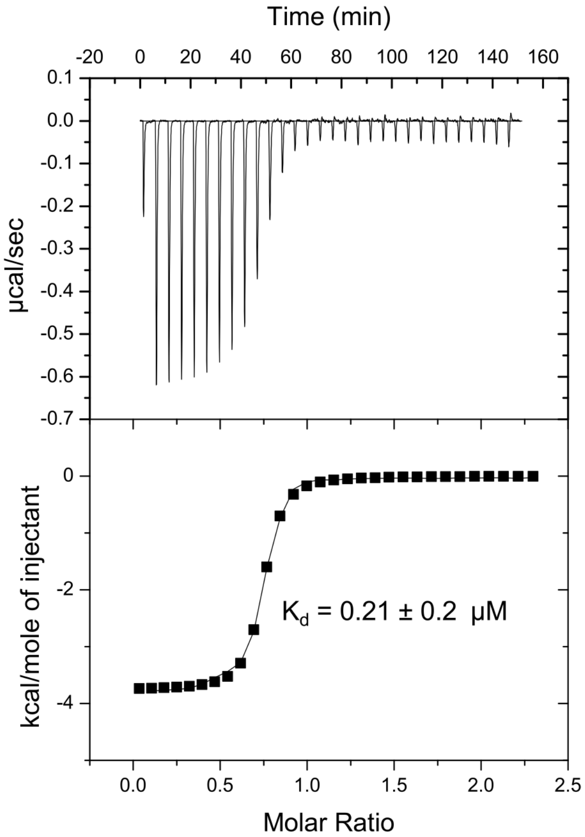


**Supplementary Figure S6.** Isothermal titration calorimetry thermogram of 800 µM TAZ2 titrated into 80 µM $E1A_{CR1}$, fit to a one-site binding model yielding $K_d$ = 0.21 ± 0.2 µM.
